# Optimisation of immunocytochemistry methodology for the detection of endogenous eIF2B localised foci

**DOI:** 10.1101/2022.12.06.519266

**Authors:** Madalena I. Ribeiro de Oliveira, Filipe M. Hanson, Rachel E. Hodgson, Alison K Cross, Susan G. Campbell, K. Elizabeth Allen

## Abstract

The multisubunit eukaryotic initiation factor 2B (eIF2B), a guanine nucleotide exchange factor (GEF) for eIF2, is an essential regulator of translation initiation. Activation of the cellular integrated stress response (ISR) by factors such as endoplasmic reticulum stress leads to phosphorylation of eIF2α and inhibition of eIF2B GEF activity. Cytoplasmic bodies containing eIF2B subunits, termed eIF2B bodies, have been shown to alter in subunit composition and fluorescence recovery after photobleaching activity in response to the ISR. Analysis of the subunit composition of endogenous eIF2B bodies is dependent on accurate detection of each protein in a cellular context via immunocytochemistry (ICC). We describe bioinformatic techniques to optimize the ICC detection of eIF2B foci in U373 cells. The screening of commercially available primary antibodies against predicted epitopes enhanced measurements of the number, size and fluorescence intensity of eIF2B bodies. A consistent and reproducible ICC analysis of endogenous eIF2B bodies will aid characterisation of eIF2B bodies during the ISR or under disease conditions.

**Summary:** eIF2B is a housekeeping protein and localised eIF2B foci, named eIF2B bodies, can be detected through immunocytochemistry. Here, we discuss the use of immunoinformatics to optimise eIF2B localisation detection.

## Introduction

The eukaryotic initiation factor 2B (eIF2B) is a guanine exchange factor (GEF) composed of five subunits (α-ε), encoded by five genes (EIF2B1-5) (Pavitt *et al*., 1998; Dev *et al*., 2010). This housekeeping protein is an essential component of translation control (Jennings & Pavitt, 2014). During translation initiation, eIF2 in its active state (eIF2-GTP) transports Met-tRNA_i_ to the small ribosomal subunit, forming the 43S preinitiation complex (PIC). Once the 43S PIC is recruited close to the mRNA m^7^G cap, it scans the mRNA and detects the AUG start codon. Upon AUG recognition, base pairing is established between the Met-tRNA_i_ anticodon and the AUG codon, eIF2-GTP is hydrolysed to eIF2-GDP and released from the PIC (Merrick & Pavitt, 2018). For subsequent rounds of translation initiation to occur, eIF2B promotes the recycling of eIF2-GDP to eIF2-GTP (Dev *et al*., 2010; Hinnebusch and Lorsch, 2012). In response to diverse cellular stresses, eIF2α is phosphorylated on Ser^51^ by a group of stress responsive kinases. This in turn, rearranges the eIF2B•eIF2 structure, inhibiting eIF2B activity, downregulating global translation and upregulating the translation of a group of stress responsive genes. This stress signalling pathway is known as the integrated stress response (ISR) and it is a critical mechanism in the pathogenesis of diverse disorders (Donnelly, Gorman, Gupta & Samali, 2013; Costa-Mattioli & Walter, 2020).

The eIF2B complex has a two-fold symmetric heterodecameric structure composed of two copies of each subunit. The subunits can be divided into subcomplexes – catalytic and regulatory. eIF2Bγ and eIF2Bε form the catalytic core, where eIF2Bε contains the nucleotide-exchange activity (Wortham *et al*., 2016). The regulatory subcomplex is composed of eIF2Bα, β and δ, which control eIF2B function during stress, by tightly binding phosphorylated eIF2α which induces a conformational change in the complex (Zyryanova, *et al*., 2021; Schoof *et al*., 2021). The regulatory subunits form a symmetric interface, with eIF2Bγ and eIF2Bε bound to the opposite ends (Wortham *et al*., 2014, 2016; Kashiwagi *et al*., 2016).

Cytoplasmic bodies, termed eIF2B bodies, are large assemblies that contain the eIF2B protein and were firstly identified in *S. cerevisiae and C. albicans* (Campbell, Hoyle and Ashe, 2005; Campbell and Ashe, 2006; Egbe *et al*., 2015; Nuske *et al*., 2020). In mammalian cells, it was found that different sized bodies are present, which vary in their subunit composition. Small bodies appear to be largely comprised of catalytic subunits, whereas medium and large bodies include the regulatory subunits (Hodgson et al., 2019). eIF2 shuttles through eIF2B bodies, and fluorescence recovery after photobleaching analysis has shown a correlation between eIF2B GEF activity and the measurement of eIF2 shuttling (Campbell, Hoyle and Ashe, 2005; Hodgson *et al* 2019; Norris *et al*., 2021). In response to stress, eIF2B subunit localisation changes occur, changing the composition of eIF2B bodies, suggesting the presence of subcomplexes which could have an important role in cellular regulation (Wortham *et al*., 2014). Furthermore, individual subcomplexes exist in different proportions, which could have an impact on stress responses and cell sensitivity (Wortham *et al*., 2014; Wortham *et al*., 2016; Hodgson *et al*., 2019).

Vanishing white matter (VWM) disease is a leukoencephalopathy that leads to neurological deterioration which is exacerbated by stress incidents such as fright or fever. VWM disease has a wide clinical spectrum, and childhood-onset is the most frequent form. It is thought to be one of the most common inherited diseases that affect white matter (van der Knaap et al., 1999; Bugiani et al., 2010). VWM is an autosomal recessive inherited disorder caused by mutations within any one of the genes encoding the five eIF2B subunits (Leegwater et al., 2001; Van Der Knaap et al., 2002; Dooves et al., 2016). Approximately 80% of the known VWM mutations are missense, with intronic variants or deletions accounting for only a small proportion (Slynko et al., 2021). The genotype-phenotype correlation is inconsistent, thus identifying how these mutations result in alterations in eIF2B localisation and subunit composition, may lead to a further understanding of the pathomechanisms of VWM disease.

To investigate the role of eIF2B bodies in the cell response to stress, immunocytochemistry (ICC) has been employed to detect and analyse eIF2B bodies in glial cells. In previous studies, GFP tagged subunits have been utilised to study the localisation of eIF2B bodies (Campbell, Hoyle and Ashe, 2005; Campbell and Ashe, 2006; Egbe *et al*., 2015; Hodgson *et al*., 2019; Nuske *et al*., 2020; Norris *et al*., 2021). However, the characterisation of endogenous complexes would allow for analysis of eIF2B bodies in patient samples and animal models of disease.

B-cell epitopes are areas of an antigen typically around five to six amino acids, which prompt an activation of immune response, binding precisely to B-cell antigen receptors. Within the structure of the Fv site of antibodies, the paratope region, i.e., the antigen-antibody site, consists of five to ten amino acids. The paratope recognizes and exclusively binds to its corresponding epitope. It is in this domain where the specific characteristics among antibodies arise (Huston, *et al*., 1996; Stave & Lindpaintner, 2013). As such, the innate proclivity that antibodies possess to bind to specific molecules allows the identification of a target protein and the observation of its distribution within cells *in situ* through techniques such as ICC (de Matos *et al*., 2010; Burry, 2010; Renshaw, 2016; Im *et al*., 2019). Reproducibility and specificity of antibodies are prominent concerns when designing and executing a particular ICC experiment. It is essential that inaccuracies concerning steps within the ICC technique are not perpetuated and that standardisation is achieved.

Bioinformatics, particularly immunoinformatics techniques have been developed for design and development of optimal antibodies, for example in reverse vaccinology (Sharma *et al*, 2022). These techniques were used recently in evaluating epitope-based vaccine design against SARS-CoV2 (Chen *et al*., 2020; Dong *et al*., 2020). Additionally, these tools can be applied to other methodologies, such as ICC, where we find molecules that are targeted by antibodies in their native form. *In silico* tools which identify optimal epitopes of proteins of interest can be used to compare commercially available antibodies or to produce custom antibodies, for reliable and optimal immunofluorescent assays.

Here we report on an immunoinformatics approach of screening the eIF2B subunit sequence and structure for antigenic and exposed areas, evaluating and selecting primary antibodies that bind to each subunit, and optimising the detection of eIF2B localised foci in glial cells. These approaches can be applicable to other complex assemblies, with or without a resolved 3D structure.

## Results

### ICC detection of endogenous eIF2B subunits in U373 cells

In previous studies, transient transfection using fluorescently tagged eIF2B subunits together with ICC methodology, have allowed the detection and analysis of eIF2B bodies (Hodgson et al., 2019). However, a comprehensive study of the distribution of endogenous subunits has yet to be carried out. To determine the localisation of the endogenous eIF2B subunits within cells, ICC analysis was carried out in U373-MG cells, utilising previously established primary antibodies to each eIF2B subunit. Antibodies were validated by western blot analysis (Figure 1A). ICC was initially carried out utilising secondary antibodies as previously published (Figure S1A and S1B) (Hodgson et al., 2019). Initial ICC analysis of endogenous eIF2B subunits showed that whilst eIF2Bα, β and δ bodies exhibited high fluorescence intensity with minimal background, eIF2Bγ and ε displayed weaker intensity (Figure S1A). Additionally, the number of eIF2B localised foci per cell differed between each subunit (Figure S1B).

**Figure 1.**
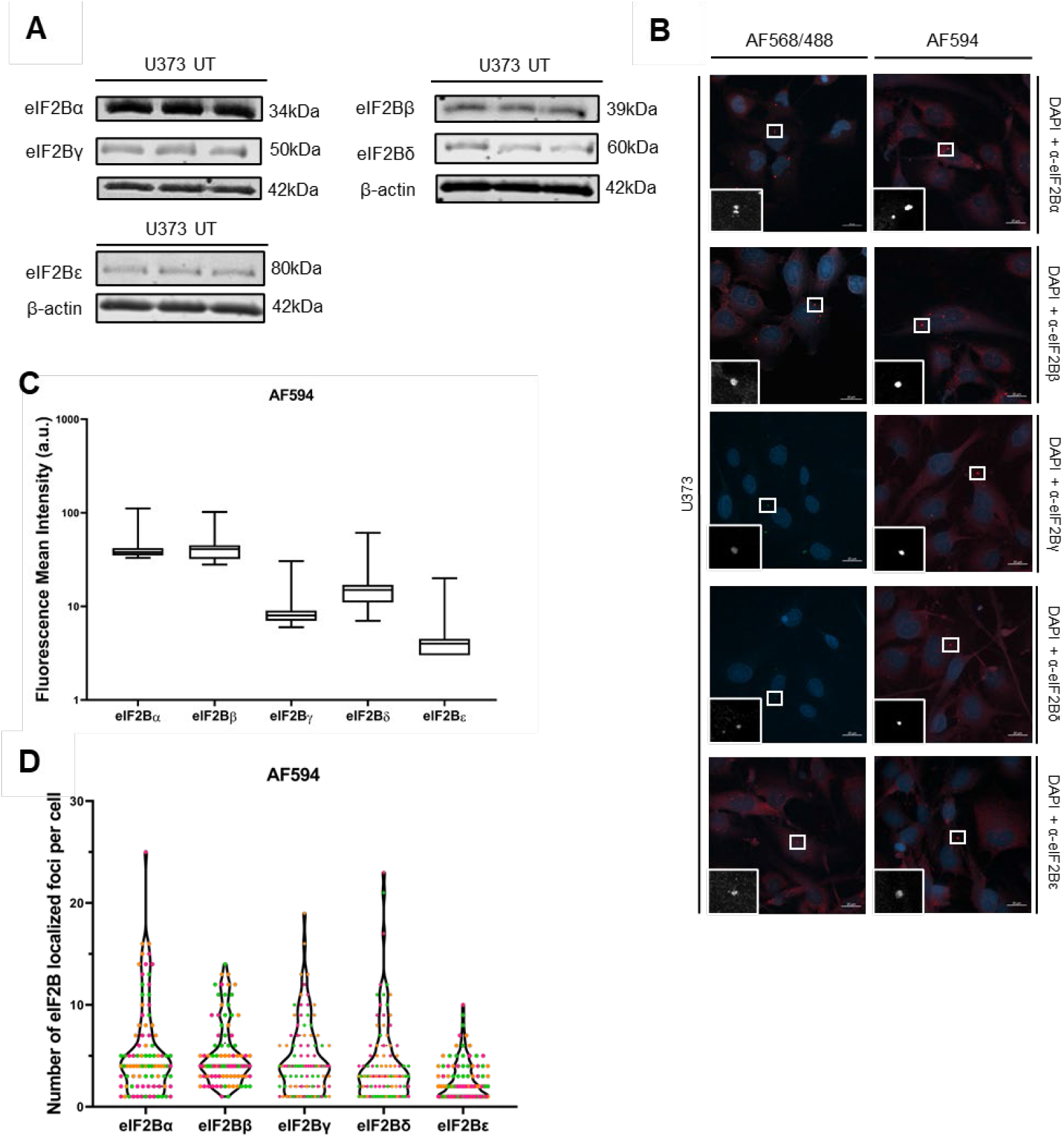
eIF2B subunits are detected in U373 cells. (A) Western Blot analysis of eIF2Bα-ε utilising previously validated primary antibodies, each lane representing a single biological replicate; (B) Confocal images of U373 cells fixed in methanol and subject to ICC with (top to bottom) primary anti-eIF2Bα, anti-eIF2Bβ, anti-eIF2Bγ, anti-eIF2Bδ and anti-eIF2Bε. Antibody staining was visualised using (left to right) Alexa Fluor 568 conjugated secondary antibody for eIF2Bα, β and ε subunits and Alexa Fluor 488 conjugated secondary antibody for eIF2Bγ and δ; appropriate Alexa Fluor 594 conjugated secondary antibodies for all subunits; (C) Fluorescence mean intensity of endogenous eIF2B subunits visualised using appropriate Alexa Fluor 594 conjugated secondary antibodies, in U373 cells (n=3 counts of 30 cells); (D) Number of eIF2B localized foci per cell detected in U373 cells visualised with appropriate Alexa Fluor 594 conjugated secondary antibodies (n=3 counts of 30 cells, n=1 in pink, n=2 in orange and n=3 in green).

In order to optimise this detection method, secondary antibodies with different fluorescent labels and antibody structure were utilised. Alpaca Fc anti-mouse conjugated to AlexaFluor 647 and alpaca Fc anti-rabbit conjugated to AlexaFluor 488 did not only fail to enhance the signal of the targeted subunits, but they also revealed weaker mean fluorescence intensity values than AlexaFluor 488 anti-mouse and AlexaFluor 568 anti-rabbit secondary antibodies. In addition Alpaca Fc secondary antibodies detected a smaller number of bodies within U373-MG cells (Figure S1C and Figure S1D). To eliminate inconsistencies between the detection of different subunits, we then utilised the same fluorescent label throughout. Anti-mouse and anti-rabbit AlexaFluor 594 conjugated secondary antibodies displayed a more consistent mean fluorescence intensity and number of localised foci per cell throughout the different eIF2B subunits (Figure 1C and 1D). Nevertheless, while Alexa Fluor 594 conjugated secondary antibodies improved overall results, the values of intensity and number of localised foci varied according to the subunit detected. eIF2Bα, β and δ generally appeared to have clear and strong signals with minimal background, while eIF2Bγ and ε displayed low intensity signals with all secondary antibodies used (Figure 1C and 1D). The varying levels of fluorescence intensity observed upon ICC analysis of specific eIF2B subunits were not related to the wavelength or structure/subtype of the secondary antibody utilised. As such, we next focused our analysis on optimising the specificity of the primary antibodies used.

### Bioinformatic analysis of the eIF2B subunits identifies viable eIF2B epitopes

In order to determine a potential source of variability in the ICC detection of endogenous eIF2B subunits and to select optimal primary antibodies, B-cell epitope prediction of each eIF2B subunit was carried out using BepiPred2.0 (Jespersen *et al*., 2017) and DiscoTope2.0 (Kringelum *et al*., 2012). These B-cell epitope analysis programmes were used to predict particularly antigenic and exposed areas for each eIF2B subunit, a methodology already utilised in other fields such as vaccine development (Chen *et al*., 2020). This allowed us to then compare these predicted antigenic sites with epitopes targeted by the primary antibodies utilised in our current ICC analysis. Exposed and antigenic sites could be altered by different conformations of the eIF2B•eIF2 complex, i.e., productive versus non-productive, or by molecules bound to the eIF2B structure (Kashiwagi *et al*., 2019; Kenner *et al*., 2019; Zyryanova, *et al*., 2018; Zyryanova, *et al*., 2021; Hao *et al*., 2021). As such, various structures with differing configurations of the eIF2B complex were used in each B-cell prediction server (Table 1).

**Table 1.** PDB IDs of eIF2B complex 3D structures.

| PDB ID | Species | Small molecules | eIF2α<br>phosphorylation<br>status | Productive/Non-productive | Reference | Notes |
| --- | --- | --- | --- | --- | --- | --- |
| **6CAJ** | Human | ISRIB | eIF2 not present | - | Tsai, *et al.*,<br>2018 | eIF2Bε catalytic domain not included |
| **6K71** | Human | - | Unphosphorylated | Productive | Kashiwagi *et al.*, 2019 | One molecule of eIF2 bound; eIF2Bε catalytic domain included |
| **6K72** | Human | - | Phosphorylated | Non-productive | Kashiwagi *et al.*, 2019 | Two molecules eIF2 bound; eIF2Bε catalytic domain not included |
| **6O9Z** | Human | ISRIB | Phosphorylated | Non-productive | Kenner *et al.*, 2019 | Two molecules of eIF2 bound, eIF2Bε catalytic domain included |
| **6O81** | Human | ISRIB | Unphosphorylated | Productive | Kenner *et al.*, 2019 | Two molecules of eIF2 bound; eIF2Bε catalytic domain included |
| **6O85** | Human | ISRIB | Unphosphorylated | Productive | Kenner *et al.*, 2019 | One molecule of eIF2 bound; eIF2Bε catalytic domain included |
| **6EZO** | Human | ISRIB | eIF2 not present | - | Zyryanova,<br>*et al.*, 2018 | eIF2Bε catalytic domain not included |
| **7D45** | Human | PHOSPHONOSERINE | Phosphorylated | Non-productive | Zyryanova,<br>*et al.*, 2021 | aP1 complex; eIF2Bε catalytic domain not included |
| **7D44** | Human | PHOSPHONOSERINE | Phosphorylated | Non-productive | Zyryanova,<br>*et al.*, 2021 | aP2 complex; eIF2Bε catalytic domain not included |
| **7D43** | Human | PHOSPHONOSERINE | Phosphorylated | Non-productive | Zyryanova,<br>*et al.*, 2021 | αPy complex, two molecules of eIF2 alpha bound; eIF2Bε catalytic domain not included |
| **7D46** | Human | PHOSPHONOSERINE | eIF2 not present | - | Zyryanova,<br>*et al.*, 2021 | eIF2Bε catalytic domain not included |

There are two main types of epitopes within a protein sequence – continuous, which are linear sequences, and discontinuous, which are present in specific complex conformation brought together by the 3D folding pattern of a protein (Baruah & Bose, 2020). Like the majority of proteins, most epitopes within the eIF2B complex are expected to be discontinuous (Barlow, Edwards & Thornton, 1986). Given the indispensable nature of the eIF2Bε subunit and the predominant number of VWM mutations within the *EIF2B5* gene, we focused our efforts on the optimization of the detection of this particular subunit. Antigenic and exposed areas of eIF2B subunits were identified using BepiPred2.0 (Figures 2A & S2) and DiscoTope2.0 (Figures 2B & S3). Amino acids highlighted by the two servers were compiled for each of the five eIF2B subunits (Figure 3A). In Figure 3B it is possible to observe an example of a viable discontinuous epitope of the eIF2Bε subunit.

**Figure 2.**
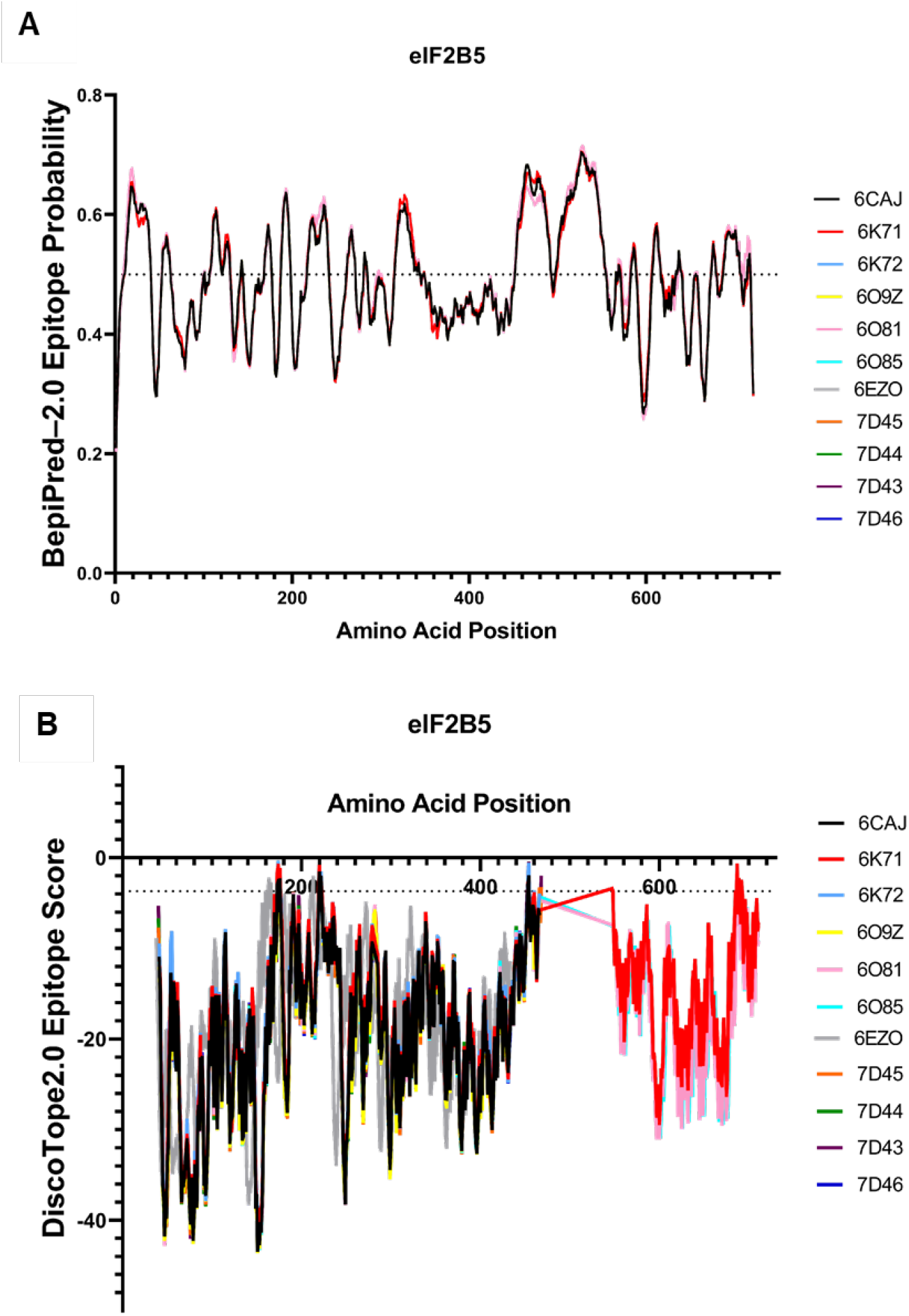
eIF2Bε epitope prediction using BepiPred–2.0 and DiscoTope2.0. (A) Bepipred-2.0 epitope probability score with 0.5 threshold of eIF2Bε using each PDB FASTA sequence; (B) DiscoTope2.0 Score with a ≥-3,7 threshold of eIF2Bε of each eIF2B PDB structure.

**Figure 3.**
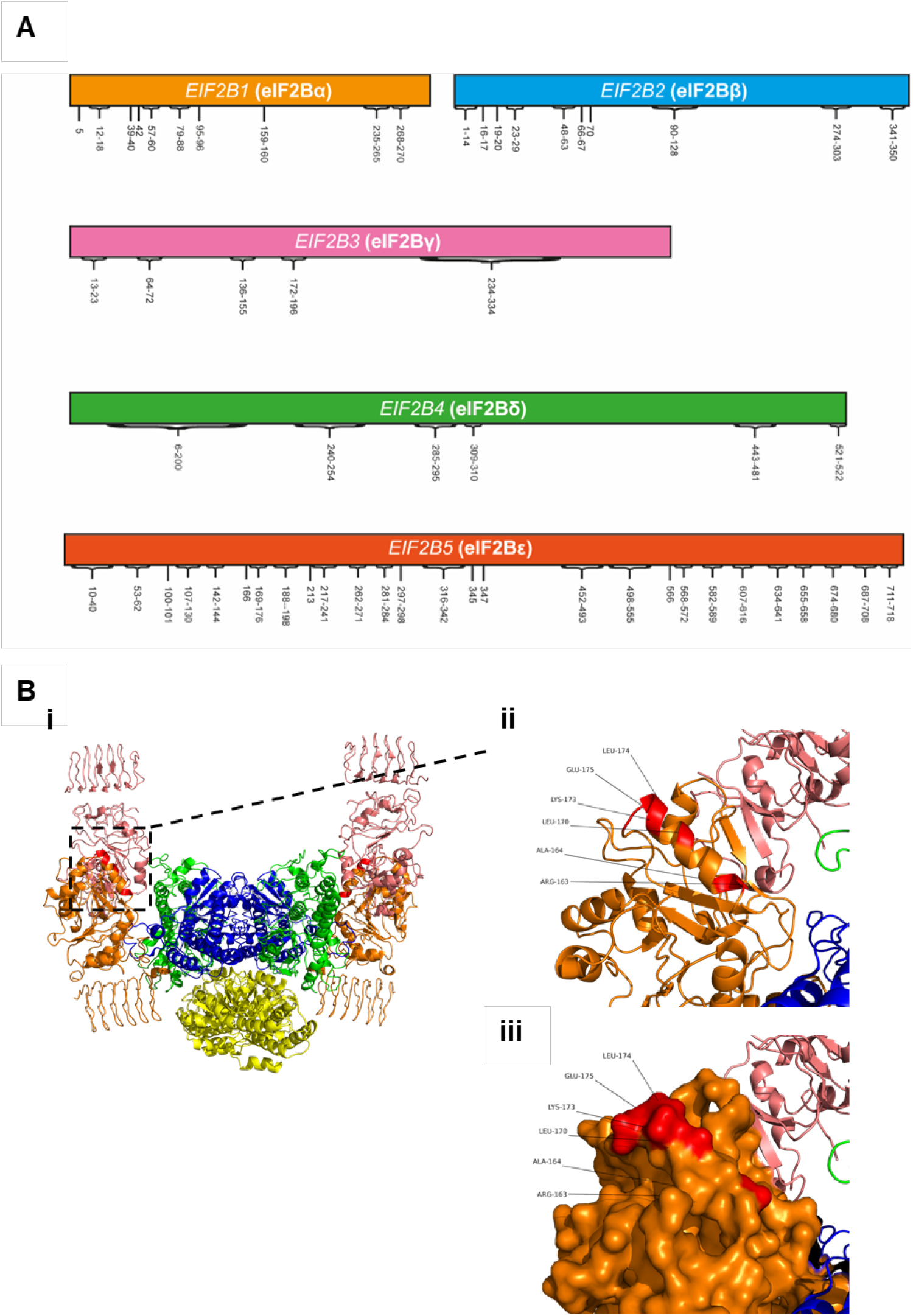
eIF2B amino acids predicted as viable epitope regions. (A) Amino acids highlighted by BepiPred-2.0 and DiscoTope2.0 servers for each eIF2B subunit. (B) Example of an eIF2Bε discontinuous epitope (PDB code: 6ezo (Zyryanova *et al*., 2018)), from the highlighted epitope in figure 3A. In each rendering, the residues that are part of the highlighted epitope are colored in red, eIF2Bα is colored yellow, eIF2Bβ is colored blue, eIF2Bγ is colored pink, eIF2Bδ is colored green, and eIF2Bε is colored orange. (i) eIF2B complex represented as ribbons. (ii) Epitope consisted of separate residues represented as ribbons; (iii) Surface rendering of the epitope.

### Predicted epitope sequences allow selection of commercially available primary antibodies targeting eIF2Bε

Following epitope analysis, a search was carried out using The Human Protein Atlas (https://www.proteinatlas.org), to identify additional primary antibodies targeting the eIF2Bε subunit. The amino acid sequence of the immunogens for the original primary antibody, ARP61329_P050, and two other commercially available primary antibodies, HPA064370 and HPA069303, were compared with the predicted epitope amino acids highlighted by the two servers (Table 2 and Figure 4A-B). Surprisingly, the initially used antibody, ARP61329_P050, displayed a higher percentage of matching amino acids with the predicted epitopes, suggesting that this antibody is optimal for the detection of the eIF2Bε subunit (Figure 4B). However, when isolating the epitopes predicted by the DiscoTope2.0 server, only HPA064370 and HPA069303 exhibited a percentage of matching amino acids, while ARP61329_P050 primary antibody did not (Table 2, Figure 4C). Figure 4D illustrates the epitopes identified by Discotope2.0 within the eIF2B decameric structure.

**Table 2.** Percentage of matching amino acids highlighted by B-cell epitope prediction servers within the paratopes of anti-eIF2Bε primary antibodies.

| Primary antibody ID | % Matching amino acids with both servers | % Matching amino acids with DiscoTope2.0 |
| --- | --- | --- |
| ARP61329_P050 | 76% | 0% |
| HPA069303 | 43% | 3% |
| HPA064370 | 4% | 2% |

**Figure 4.**
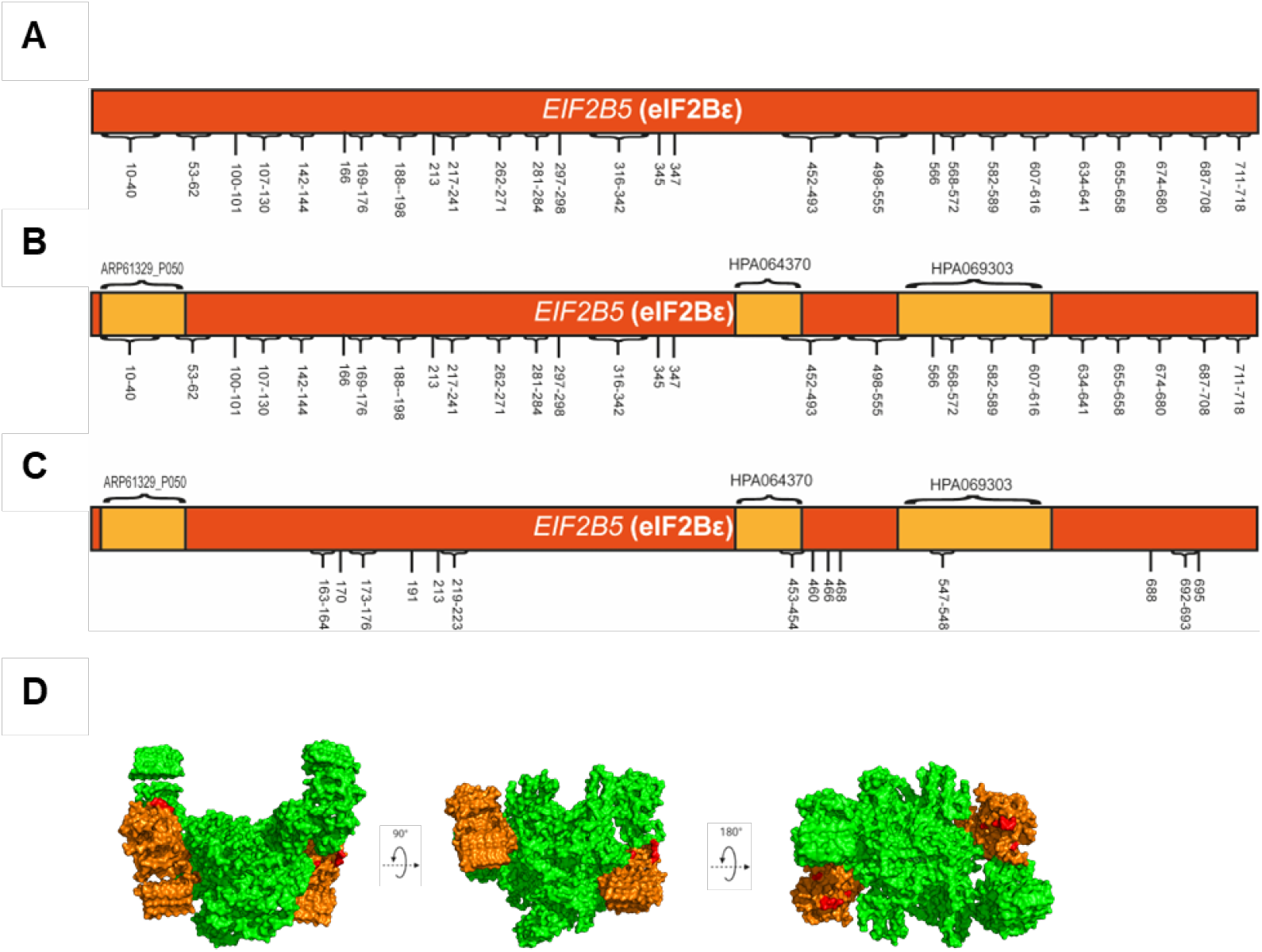
eIF2Bε amino acids predicted as viable epitope regions. (A) Amino acids highlighted by both of the epitope prediction servers (BepiPred-2.0 and DiscoTope2.0) for the eIF2Bε subunit; (B) Amino acids highlighted by both of the epitope prediction servers (BepiPred-2.0 and DiscoTope2.0) for the eIF2Bε subunit and the binding sites of primary antibodies that target this protein; (C) Amino acids highlighted by DiscoTope2.0 for the eIF2Bε subunit and the binding sites of primary antibodies that target this protein; (D) PyMOL imaging of DiscoTope2.0 highlighted eIF2Bε epitopes. Green – eIF2B complex; yellow – eIF2Bε; red – Highlighted epitopes (PDB code: 6ezo (Zyryanova *et al*., 2018)).

To determine if the HPA064370 and HPA069303 antibodies displayed a higher fluorescence intensity, ICCs using these antibodies were carried out (Figure 5A). An increase of mean fluorescence intensity was shown with HPA064370 and HPA069303 primary antibodies when compared with ARP61329_P050, the original antibody used (Figure 5B). Additionally, both HPA064370 and HPA069303 primary antibodies identified a greater number of eIF2B localised foci per cell, closer to the values obtained for the other eIF2B subunits (Figure 5C). We further analysed the detection of endogenous eIF2Bε with and without endoplasmic reticulum stress induced by thapsigargin, to determine whether the detection of eIF2Bε was altered by conformational changes caused by binding of phosphorylated eIF2α. The number of eIF2Bε bodies detected with all three primary antibodies increased upon thapsigargin treatment (Fig. 5C). This increase in body number was not due to a change in the protein expression levels of eIF2Bε (Fig S4).

**Figure 5.**
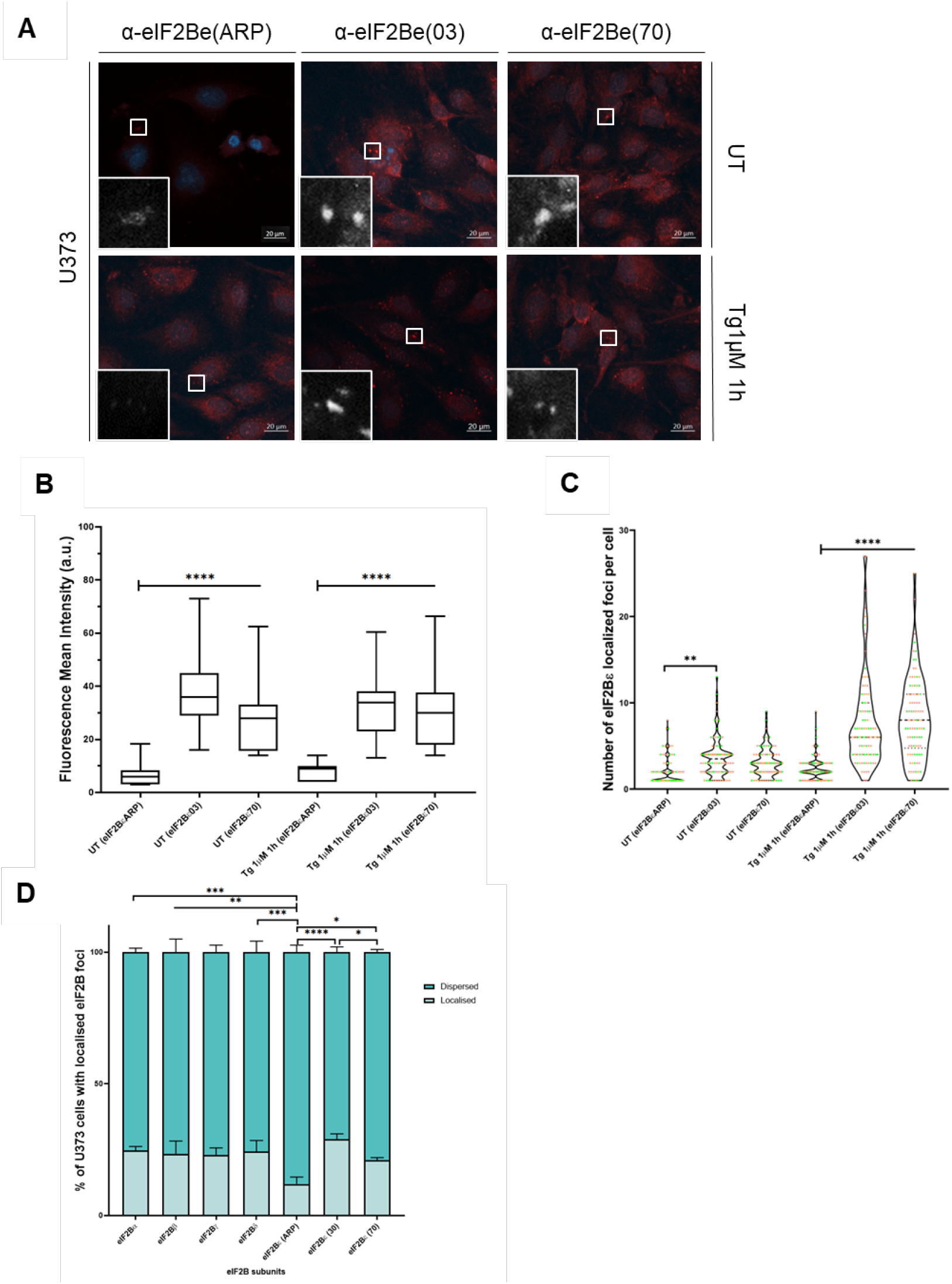
Immunoinformatics optimisation of primary antibodies results in an increase in mean fluorescence intensity of eIF2Bε localization. (A) Confocal images of endogenous eIF2Bε localized foci in U373 cells. Cells were treated with vehicle or 1μM thapsigargin for one hour; (B) Fluorescence mean intensity analysis of U373 cells subject to ICC with three primary antibodies to the eIF2Bε subunit. Cells were treated with vehicle or 1μM thapsigargin for one hour (n=3, counts of 30 cells). p values were derived from a Kruskal-Wallis test, followed by a Conover-Inman analysis, * * * *p ≤0.0001; (C) Number of eIF2Bε localization detected in a population of U373 cells with ARP61329_P050, HPA069303 and HPA064370 primary antibodies targeting eIF2Bε. Cells were treated with vehicle or 1μM thapsigargin for one hour (n=3 counts of 30 cells, n=1 in pink, n=2 in orange and n=3 in green). p values were derived from a Kruskal-Wallis test, followed by a Conover-Inman analysis, * * *p ≤0.001 and * * * *p ≤ 0.0001; (D) Percentage of U373 cells with eIF2Bα-ε localization (n=3, counts of 100 cells)

Following the optimisation of the primary antibody targeting the eIF2Bε subunit of the eIF2B protein, the characterisation of endogenous eIF2B bodies detected by antibodies to each individual subunit was carried out in U373 cells. The difference in percentage of cells with localised foci was statistically significant between the three primary antibodies to eIF2Bε (Figure 5D). The percentage of cells with eIF2B localised foci, was consistent for ICC analysis of each eIF2B subunit, when either HPA064370 and HPA069303 primary antibodies were utilised to detect eIF2Bε (Figure 5D).

## Discussion

The eIF2B protein complex localises to cytoplasmic foci, termed eIF2B bodies. These bodies in mammalian cells have distinct sizes, which correlate with subunit composition (Hodgson *et al*., 2019). VWM disease is an autosomal recessive inherited chronic-progressive neurodegenerative disorder caused by mutations within the genes encoding the five eIF2B subunits (van der Knaap *et al*., 2002). Given that mutations in the eIF2B subunits associated with VWM do not always affect the biochemical activity of eIF2B (Liu *et al*., 2011), understanding the localisation pattern of eIF2B bodies may uncover possible undetermined eIF2B functionalities within cells and obtain a greater insight into the correlation of genotype-phenotype of VWM mutations and cell specificity of the causative VWM mutations. Here we present data on the optimization of ICC analysis of endogenous eIF2B localisation foci and their characterisation in a glial cell line. Previous characterisation of eIF2B bodies was carried out utilising exogenously expressed GFP tagged proteins (Campbell, Hoyle and Ashe, 2005; Campbell and Ashe, 2006; Egbe *et al*., 2015; Hodgson *et al*., 2019; Nuske *et al*., 2020; Norris *et al*., 2021). We aimed to detect endogenous eIF2B subunits in U373-MG cells to allow evaluation of endogenous mutant proteins within patient cells or animal models.

Initially, ICC analysis was carried out using formerly published primary and secondary antibodies (Hodgson *et al*., 2019). The endogenous eIF2B subunits were detected (Figure 1B); however, an inconsistency in the detected signal intensity and number of localised foci across the subunits was observed (Figure S1A and S1B). The initial optimisation focused on the secondary antibodies as the primary antibodies used had been previously validated (Hodgson et al., 2019). We postulated that by using alpaca Fc anti-mouse conjugated AlphaFluor 647 and alpaca Fc anti-rabbit conjugated AlphaFluor 488 secondary antibodies we would obtain better results, given they are smaller antibodies, providing a better epitope access (*The ChromoTek Alpaca Antibody Advantage* | *Chromotek*, n.d.). However, low levels of fluorescent intensity and a decrease in the number of eIF2B bodies were observed (Figure S1C, S1D and S1E). Alexa Fluor anti-mouse and anti-rabbit 594 were chosen for subsequent ICC analysis to detect all eIF2B localised foci, due to exhibiting a more consistent signal intensity. Additionally, these antibodies have identical excitation wavelengths thereby eliminating this particular detection variability. These secondary antibodies displayed improved signals, however antibodies targeting the eIF2Bγ and ε subunits displayed weaker intensity throughout the range of secondary antibodies (Figure 1C). Additionally, a lower number of localised eIF2Bε foci was observed when compared to other eIF2B subunits (Figure 1D). When comparing the different mean fluorescence intensity levels of endogenous eIF2B bodies, the results were not dependent on the antibody species, since the subunits with the lower intensity are seen with both anti-mouse and anti-rabbit secondary antibodies. This variability in body number could be due to the localisation pattern of eIF2B subunits within U373-MG cells, or due to low intensity signals. While assessing foci localisation, it is crucial to have a fluorescence intensity signal higher than background. Therefore, experimental approaches to eliminate poor antibody detection as a limiting factor for the reliable ICC characterisation of these cytoplasmic foci were carried out.

The primary antibodies used for the ICC methodology were examined as a possible cause for the discrepancies in fluorescence intensity and number of foci detected for the individual subunits. When choosing a primary antibody, it is important to consider the methodology being employed. Western Blot analysis utilises antibodies to target the linear structure of unfolded proteins. On the other hand, ICC analysis employs primary antibodies that target proteins with their nearly intact native conformation (non-denatured) form (Bordeaux *et al*., 2010). As such, the surface accessibility of targeted epitopes is an important topic to consider when choosing primary antibodies for ICC. This is particularly applicable for the eIF2B complex, which has been shown to alter its conformation in response to interaction with molecules such as phosphorylated eIF2α (eIF2(α-P)) or the ISR modulator ISRIB (reviewed in Marintchev & Ito, 2020; Zyryanova *et al*., 2021; Schoof *et al*., 2021). The complex structure of eIF2B•eIF2 can be found in two states – productive and non-productive. Following binding of eIF2(α-P) to eIF2B, the conformation of the eIF2B βδγε tetrameric unit shifts to a non-productive state, which antagonises catalytically productive binding of eIF2 to the β and δ subunits of eIF2B (Zyryanova *et al*., 2021; Schoof *et al*., 2021). This conformational change within the eIF2B decamer may in turn change the epitope accessibility. An array of human eIF2B structures were analysed, representing the productive (eIF2B•eIF2) and non-productive (eIF2B•eIF2(α-P)), structures of the eIF2B complex. Additionally, structures bound to ISRIB have been analysed (Table 1).

To determine viable and exposed epitopes of the eIF2B subunits, two distinct B-cell epitope servers were used. BepiPred – 2.0 is a linear B-cell epitope prediction programme that uses epitope information obtained from a protein sequence to examine the physico-chemical characteristics utilising a Random Forest algorithm (Jespersen *et al*., 2017) (Figure 2A and S2). DiscoTope 2.0 discontinuous B-cell epitope prediction server uses the 3D structure information of a protein. This server analyses the contact numbers, which is the spatial distance between residue-residue of 3D structures. This in turn determines the surface accessibility and propensity scores of amino acid residues according to the selected threshold for epitope identification, being the default value of -3.7 (Kringelum *et al*., 2012) (Figure 2B and S3). Additionally, there are various other prediction programmes that apply similar prediction methods, focusing on linear epitopes, such as ABCpred (Saha & Raghava, 2006) or on conformational (discontinuous) epitopes, such as ElliPro (Ponomarenko *et al*., 2008.).

Amino acids highlighted by the two servers for each eIF2B subunit were determined (Figure 3). The immunogens used to generate eIF2Bα and β antibodies were EIF2B1 and EIF2B2 full sequence fusion proteins, respectively, therefore information of potential epitopes detected by these antibodies was not available. We then looked at the eIF2Bε analysis in more detail. The regions of the eIF2Bε protein used to raise the primary antibodies ARP61329_P050, HPA069303 and HPA064370 were compared to the bioinformatic analysis of epitopes within the eIF2Bε subunit (Table 2 and Figure 4). From this analysis, it was found that ARP61329_P050 had the highest percentage match between the region used to raise the antibody and antigenic regions within the eIF2Bε protein (Table 2 and Figure 4B).

Epitopes can be divided into continuous – linear amino acid sequences; and discontinuous - amino acid sequences with a folded conformation, which are the large majority of epitopes (Moreau, Granier, Villard, Laune & Molina, 2006). As DiscoTope2.0 is the only server utilised that identifies discontinuous epitopes, we evaluated its results alone (Table 2, Figure 4C and 4D). Whilst DiscoTope 2.0 highlighted the lowest number of antigenic amino acids throughout the five eIF2B subunits (Figure 2B and S3), eIF2Bε peptides used to raise HPA064370 and HPA069303 antibodies appeared to match with antigenic regions identified by DiscoTope analysis, while ARP61329_P050 did not. This suggests that the ARP61329_P050 antibody has an antigenic region on the linear protein, however within the native eIF2B epsilon structure this linear region is not antigenic. It is important to note that the number of highlighted amino acids using DiscoTope2.0 was scarce throughout the subunits, but particularly in eIF2Bε.

To determine if these bioinformatic results would correlate with experimental ICC detection of eIF2Bε, we analysed the fluorescence intensity and number of localised eIF2Bε foci using ARP61329_P050, HPA064370 and HPA069303 primary anti-eIF2Bε antibodies, detected with the appropriate secondary antibody conjugated to Alexa Fluor 594. We found that HPA064370 and HPA069303 antibodies significantly increased the mean fluorescent signal intensity of the endogenous eIF2Bε foci (Figure 5B). These results support the proposed experimental approach of utilising B-cell epitope servers for optimisation of ICC detection of proteins, and indicates that the epitope identified by DiscoTope2.0 is more accessible. We suggest that this approach is a useful tool to select commercially available primary antibodies for ICC techniques, which would allow for troubleshooting of suboptimal ICC results. Additionally, immunoinformatics enables for the design of custom primary antibodies with optimal antigenicity, which has been reported previously (Chen *et al*., 2020; Liew *et al*., 2021).

Binding of Ser^51^ phosphorylated eIF2α alters the conformation of the eIF2B complex (Zyryanova *et al*., 2021; Schoof *et al*., 2021). Our data show that the immunoinformatics analysis does not detect epitope changes between the various 3D structures of the eIF2B•eIF2 complex (Figures 2, S2 & S3). This is unsurprising given the 1-6 Å scale of conformational changes within the eIF2B complex upon binding of eIF2(α-P) or ISRIB (Zyryanova *et al*., 2021; Schoof *et al*., 2021). We then went on to analyse the detection of endogenous eIF2Bε with and without endoplasmic reticulum stress, to determine whether the detection of eIF2Bε was altered by conformational changes. The number of epsilon bodies detected with all three primary antibodies increased upon thapsigargin treatment (Fig. 5A & 5C). This increase is comparable for all three anti-eIF2Bε primary antibodies, suggesting that it is an increase of epsilon body formation rather than a change in epsilon conformation that has been detected. This increase in body number was not due to a change in the protein expression levels of eIF2Bε (Fig S4).

Immunoinformatics analysis of protein epitopes and comparison of subunit – subunit interfaces within the eIF2B complex could be used to develop molecular probes to investigate the formation of eIF2B subcomplexes within a cellular context, an approach already utilised in other fields (de Beer & Giepmans, 2020) Following the optimisation of ICC detection of eIF2Bε, analysis of all eIF2B subunits in U373-MG cells was carried out. The analysis showed that the primary antibody utilised has a statistically significant impact on the percentage of cells identified to contain eIF2B bodies (Fig. 5D). For future studies, the reliable analysis and comparison between neuronal and glial cells will allow for the characterisation of eIF2B body prevalence in a cell-type specific manner. Additionally, colocalization of different eIF2B subunits will establish the correlation between body size and subunit composition in different cell lines.

To conclude, ICC is a powerful methodology to detect the cellular localisation of proteins *in situ*, yet the selection of primary antibodies used is often overlooked. We here describe bioinformatic tools that can aid antibody screening and selection, thus facilitating the study of non-denatured complex structures, such as eIF2B. This allows for inexpensive optimization, reliable detection and analysis of proteins. However, it is important to consider the types of B-cell epitope prediction servers employed - sequence or structured based - and the protein complex used for analysis – bound to other molecules or not, large portions of unmodeled sections – which could translate into limiting factors for the bioinformatic analysis, impacting the overall results.

## Materials and methods

### Cell culture

Human U373-MG astrocytoma cells were cultured in Mininum Esssential Medium (MEM), supplemented with 10% (vol/vol) fetal bovine serum (FBS), 1% (wt/vol) glutamine, 1% (wt/vol) non-essential amino acids, 1% (wt/vol) penicillin/streptomycin and 1% (wt/vol) sodium pyruvate. Cells were maintained at 37ºC under 5% CO_2_ in a humidified atmosphere and routinely tested for contamination with MycoAlter™ Mycoplasma Detection Kit (Lonza, UK). Cells were validated through lineage-specific markers - anti-GFAP antibody, a gift from Prof Woodroofe (Sheffield Hallam University). All experiments were carried out with cellular passage number no higher than 23. All media and supplements were purchased from Life Technologies Co. (New York, USA), except when indicated otherwise.

### Cell treatments

For induction of cellular stress, U373-MG cells were treated with 1 μM thapsigargin (Tg) (stock solution: 1 mg/mL diluted in DMSO stored at -20°C; Sigma-Aldrich) for 60 minutes at 37°C.

### Western Blot analysis

For protein extraction, cells were washed with PBS (Sigma-Aldrich) and lysed in in CelLytic M (Sigma-Aldrich) freshly supplemented for each use with 10mM sodium fluoride (NaF), 1mM PMSF, 17.5 mM β-glycerophosphatase, 1% (v/v) phosphatase inhibitor cocktail 2 (Sigma-Aldrich), 1% (v/v) phosphatase inhibitor cocktail 3 (Sigma-Aldrich) and 1% (v/v) protease inhibitor cocktail (Sigma-Aldrich) for 15 min at 4°C with regular agitation. Cell lysates were centrifugated (13,000 rpm for 10 minutes at 4°C) and protein concentration was determined via Qubit™ Protein Assay Kit (Thermo-Fisher), following manufacturer’s instructions.

Protein samples were diluted in 4x SDS-PAGE sample buffer supplemented with fresh β-mercaptoethanol (10% (vol/vol)) and incubated at 95ºC for 5 min. Total protein (20 μg) was separated through a 10 % polyacrylamide gel. 1-2 μL of Chameleon Duo Pre-Stained Protein Ladder (LiCor) was used as a molecular weight marker. Gels were wet-transferred onto nitrocellulose membrane (Amersham™ Protran®; Sigma-Aldrich). Membranes were blocked in 5 % (wt/vol) non-fat milk or BSA in TBST and probed with primary antibodies diluted in TBST supplemented with 5 % (wt/vol) non-fat milk or BSA. The primary antibodies used are displayed in Table S1. Membranes were then washed with TBST, probed with LiCor secondary antibodies and visualised on a LiCor Odyssey Scanner with Image Studio Lite software. The secondary antibodies used are displayed in Table S1.

### Immunocytochemistry

Cells were grown on coverslips in six-well plates. The cells were fixed in ice-cold methanol at -20ºC for 15 min and then washed with PBS supplemented with 0.5% (vol/vol) Tween 20 (PBST), three times for 5 min. Finally, the cells were blocked in 1% (wt/vol) BSA (diluted in PBS). Following blocking, cells were washed with PBST, three times for 5 min, and incubated with primary antibodies in 1% (wt/vol) BSA (diluted in PBS) overnight at 4ºC. The primary antibodies used are displayed in Table S1. Cells were washed with PBST, four times for 5 min, and then incubated with the appropriate secondary antibody (Thermo Fisher Scientific 1% (wt/vol) BSA (diluted in PBS) for 1h at RT. The secondary antibodies used are displayed in Table S1. Cells were then washed with PBST 4 times for 5 min and mounted using ProLong™ Gold Antifade Mountant with DAPI 220 (Invitrogen, Thermo Fisher Scientific). Cells were viewed on a Zeiss LSM 800 confocal microscope.

### Confocal Imaging

Imaging was performed using Zeiss LSM 800 confocal microscope with Zeiss ZEN 2.3 (blue edition) software for data processing and analysis. A 63X plan-apochromat oil objective and a diode laser with a maximum output of 1% laser transmission was used for LSM confocal imaging. Orthogonal projection of Z-stack images of automatically calculated increments were used for image acquisition. To identify eIF2B foci, cell images were processed for automatic detection for respective fluorescence prior manual setup of intensity threshold.

### Prediction of B-cell epitopes

B-cell epitope prediction was performed with the use of BepiPred2.0 (Jespersen *et al*., 2017) and DiscoTope2.0 (Kringelum *et al*., 2012) servers. PyMOL was used as a visualisation tool of the eIF2B complex and the localization of epitopes of primary antibodies that bind to the complex (Rigsby & Parker, 2016).

### Measurement and statistical analysis

To determine the percentage of matching amino acids highlighted by B-cell epitope prediction servers with the paratopes of primary antibodies, the sequence of each eIF2B subunit was thought as 100% and the total number of the amino acids highlighted was compared in proportion with the total sequence. Data was subjected to Shapiro-Wilk normality test. If parametric, data was analysed by one-way ANOVA test for comparison of three of more groups followed by Tukey’s correction post-hoc test. For analysis of groups split on two variables two-way ANOVA test, followed by a Sidak’s analysis was carried out. Data were considered parametric when *p* < 0.05. Asterisks indicate respective statistical significance as follows: \**p* < 0.05; \*\**p* < 0.01; \*\*\**p* < 0.001; and \*\*\*\**p* < 0.0001.

---

**Supplementary Table S1.**

| Primary antibodies for Western Blot Analysis |  |  |  |
| --- | --- | --- | --- |
| Antibody Target | Catalogue Number | Supplier | Dilution |
| β-actin | #3700 | Cell Signalling | 1:2500 |
| GAPDH | #2118 | Cell Signalling | 1:5000 |
| eIF2Bα | 18010-1-AP | Proteintech | 1:500 |
| eIF2Bβ | 11034-1-AP | Proteintech | 1:500 |
| eIF2Bγ | sc-137248 | Santa Cruz Biotechnology | 1:500 |
| eIF2Bδ | sc-271332 | Santa Cruz Biotechnology | 1:500 |
| eIF2Bε | ARP613229-P050 | Aviva Systems Biology | 1:500 |
| eIF2Bε | HPA064370 | Sigma-Aldrich | 1:500 |
| eIF2Bε | HPA069303 | Sigma-Aldrich | 1:500 |
| eIF2α | ab5369 | Abcam | 1:500 |
| phosho-eIF2α[ser51] | ab32157 | Abcam | 1:500 |
| Secondary antibodies for Western Blot Analysis |  |  |  |
| Antibody Target | Catalogue Number | Supplier | Dilution |

| Goat-anti-mouse IRDye 680RD | 925-68070 | LiCor | 1:10000 |
| --- | --- | --- | --- |
| Goat-anti-rabbit IRDye 800CW | 926-32211 | LiCor | 1:10000 |
| Primary antibodies for Immunocytochemistry Analysis |  |  |  |
| Antibody Target | Catalogue Number | Supplier | Dilution |
| eIF2Bα | 18010-1-AP | Proteintech | 1:25 |
| eIF2Bβ | 11034-1-AP | Proteintech | 1:25 |
| eIF2Bγ | sc-137248 | Santa Cruz<br>Biotechnology | 1:50 |
| eIF2Bδ | sc-271332 | Santa Cruz<br>Biotechnology | 1:50 |
| eIF2Bε | ARP61329_P050 | Aviva Systems<br>Biology | 1:500 |
| eIF2Bε | HPA064370 | Sigma-Aldrich | 1:100 |
| eIF2Bε | HPA069303 | Sigma-Aldrich | 1:25 |
| Secondary antibodies for Immunocytochemistry Analysis |  |  |  |
| Antibody Target | Catalogue Number | Supplier | Dilution |
| AlexaFluor 488 anti-mouse | A28175 | Thermo Fisher | 1:500 |
| AlexaFluor 568 anti-rabbit | A-11011 | Thermo Fisher | 1:500 |
| Alpaca anti-rabbit, Alexa Fluor<br>488 | srbAF488-1-10 | ChromoTek | 1:500 |
| Alpaca anti-mouse, Alexa Fluor<br>647 | sms1AF647-1-10 | ChromoTek | 1:500 |
| AlexaFluor 594 anti-rabbit | A-11012 | Thermo Fisher | 1:500 |
| AlexaFluor 594 anti-mouse | R37121 | Thermo Fisher | 1:500 |

## Acknowledgements

We thank our colleague Dr Celine Soulihol for her technical expertise and advice. For the purpose of open access, the author has applied a Creative Commons Attribution (CC BY) licence to any Author Accepted Manuscript version arising from this submission.

## Data availability

All data are included within the article.

## Figures & legends

**Figure S1.**
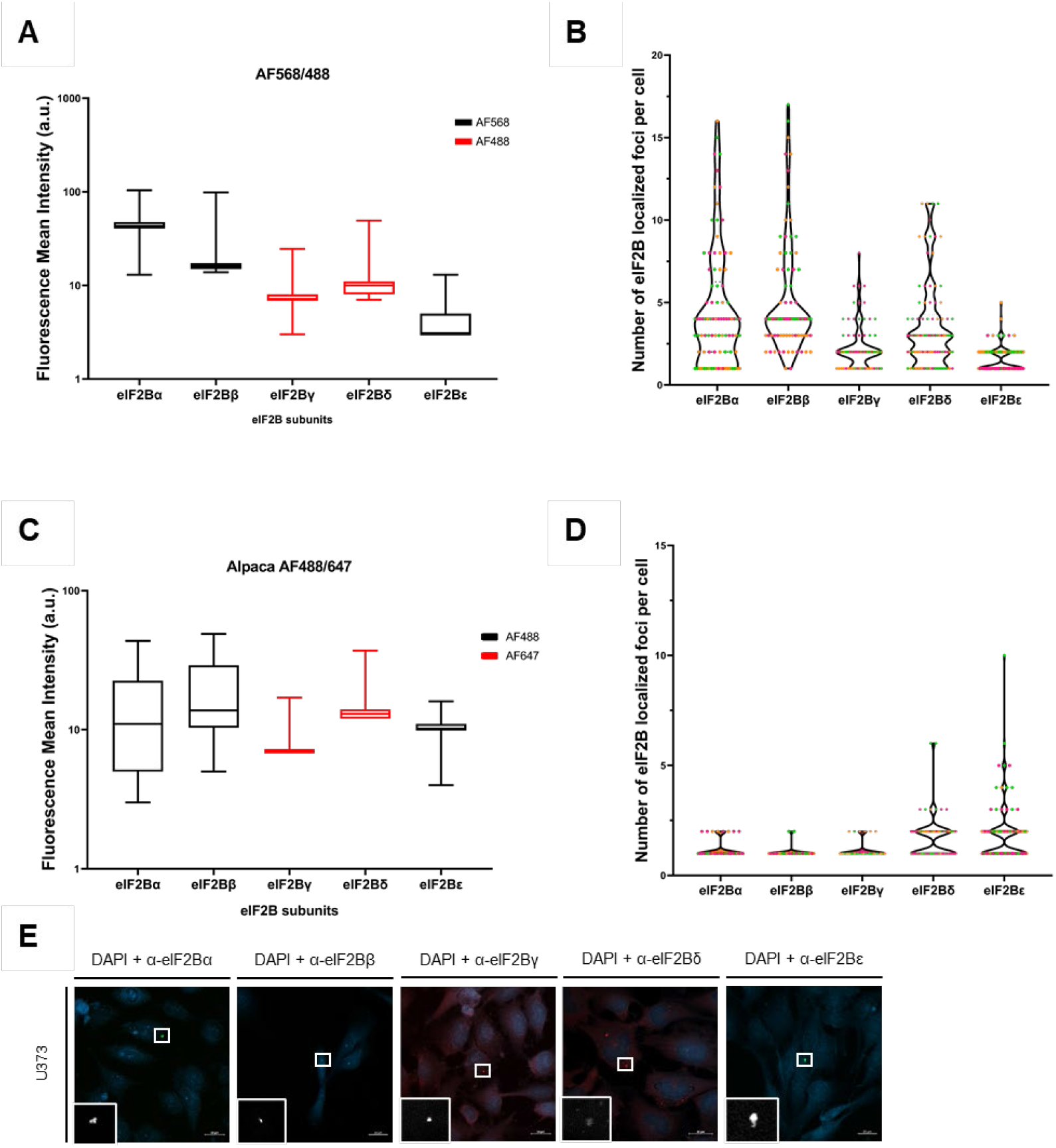
eIF2Bγ and ε subunits have low mean fluorescence intensity values throughout different secondary antibodies. (A) Fluorescent mean intensity of endogenous eIF2B subunits visualised using Alexa Fluor anti-rabbit 568 conjugated secondary antibody for eIF2Bα, β and ε subunits and Alexa Fluor anti-mouse 488 conjugated secondary antibody for eIF2Bγ and δ. Individual eIF2B localized foci were selected within U373 cells and ZEN 2.3 (blue edition) software was used for arithmetic mean intensity analysis. (n=3, counts of 30 cells); (B) Number of eIF2B localized foci per cell detected in U373 cells with Alexa Fluor anti-rabbit 568 conjugated secondary antibody for eIF2Bα, β and ε subunits and Alexa Fluor anti-mouse 488 conjugated secondary antibody for eIF2Bγ and δ. (n=3 counts of 30 cells, n=1 in pink, n=2 in orange and n=3 in green); (C) Fluorescent mean intensity of endogenous eIF2B subunits visualised using Alpaca Fc anti-rabbit 568 conjugated secondary antibody for eIF2Bα, β and ε subunits and Alpaca Fc anti-mouse 488 conjugated secondary antibody for eIF2Bγ and δ. Individual eIF2B localized foci were selected within at least 10 U373 cells and ZEN 2.3 (blue edition) software was used for arithmetic mean intensity analysis. (n=3, counts of at least 10 cells); (D) Number of eIF2B localized foci per cell detected in U373 cells using Alpaca Fc anti-rabbit 568 conjugated secondary antibody for eIF2Bα, β and ε subunits and Alpaca Fc anti-mouse 488 conjugated secondary antibody for eIF2Bγ and δ, (n=3, counts of at least 10 cells, n=1 in pink, n=2 in orange and n=3 in green); (E) Confocal images of U373 cells fixed in methanol and subject to ICC with (left to right) primary anti-eIF2Bα, anti-eIF2Bβ, anti-eIF2Bγ, anti-eIF2Bδ and anti-eIF2Bε. Antibody staining was visualised using Alpaca Fc anti-rabbit 568 conjugated secondary antibody for eIF2Bα, β and ε subunits and Alpaca Fc anti-mouse 488 conjugated secondary antibody for eIF2Bγ and δ.

**Figure S2.**
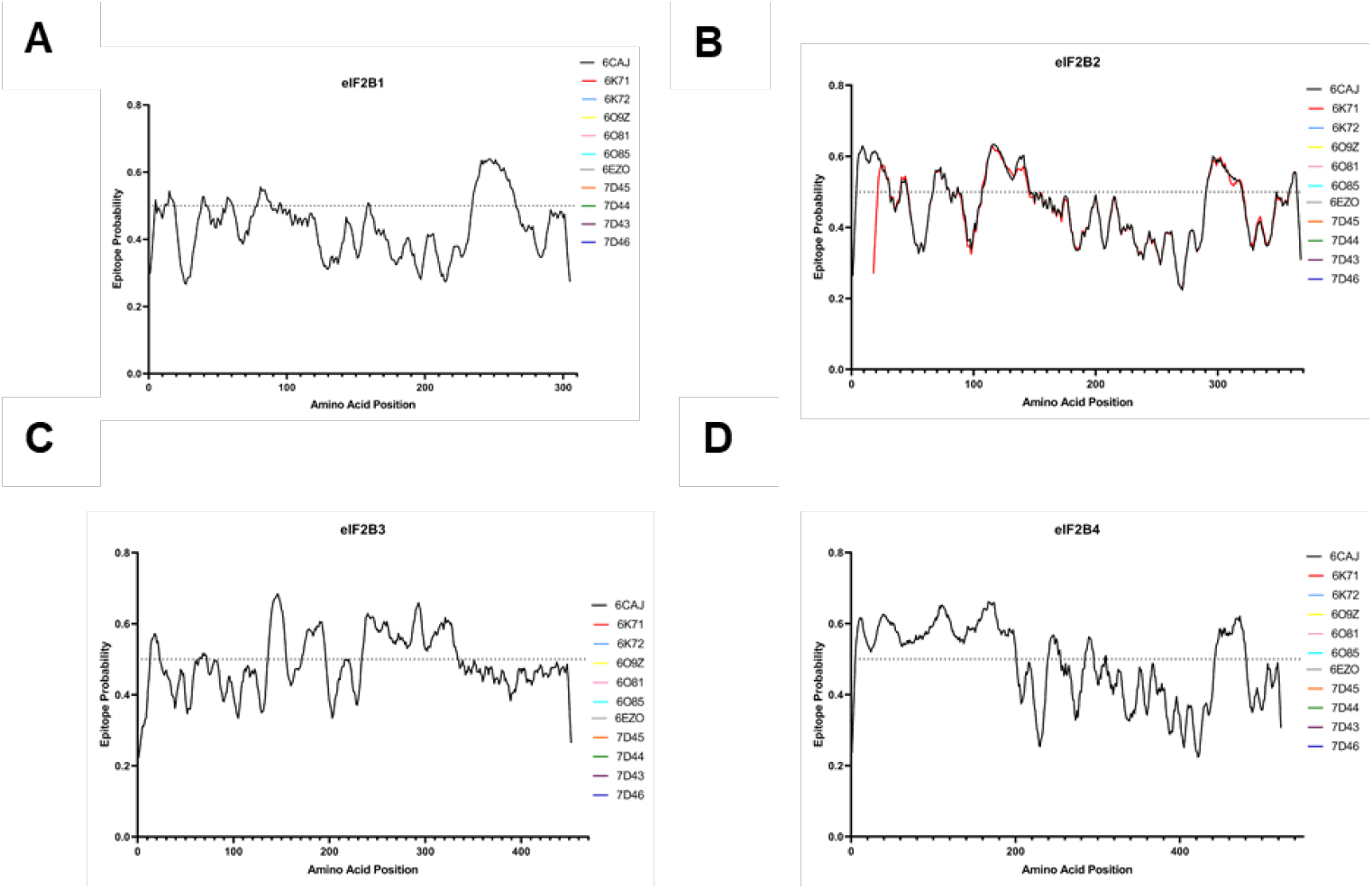
eIF2B epitope prediction using BepiPred–2.0. (A) Epitope probability score with 0.5 threshold of eIF2Bα using each PDB FASTA sequence; (B) Epitope probability score with 0.5 threshold of eIF2Bβ using each PDB FASTA sequence; (C) Epitope probability score with 0.5 threshold of eIF2Bγ using each PDB FASTA sequence; (D) Epitope probability score with 0.5 threshold of eIF2Bδ using each PDB FASTA sequence.

**Figure S3.**
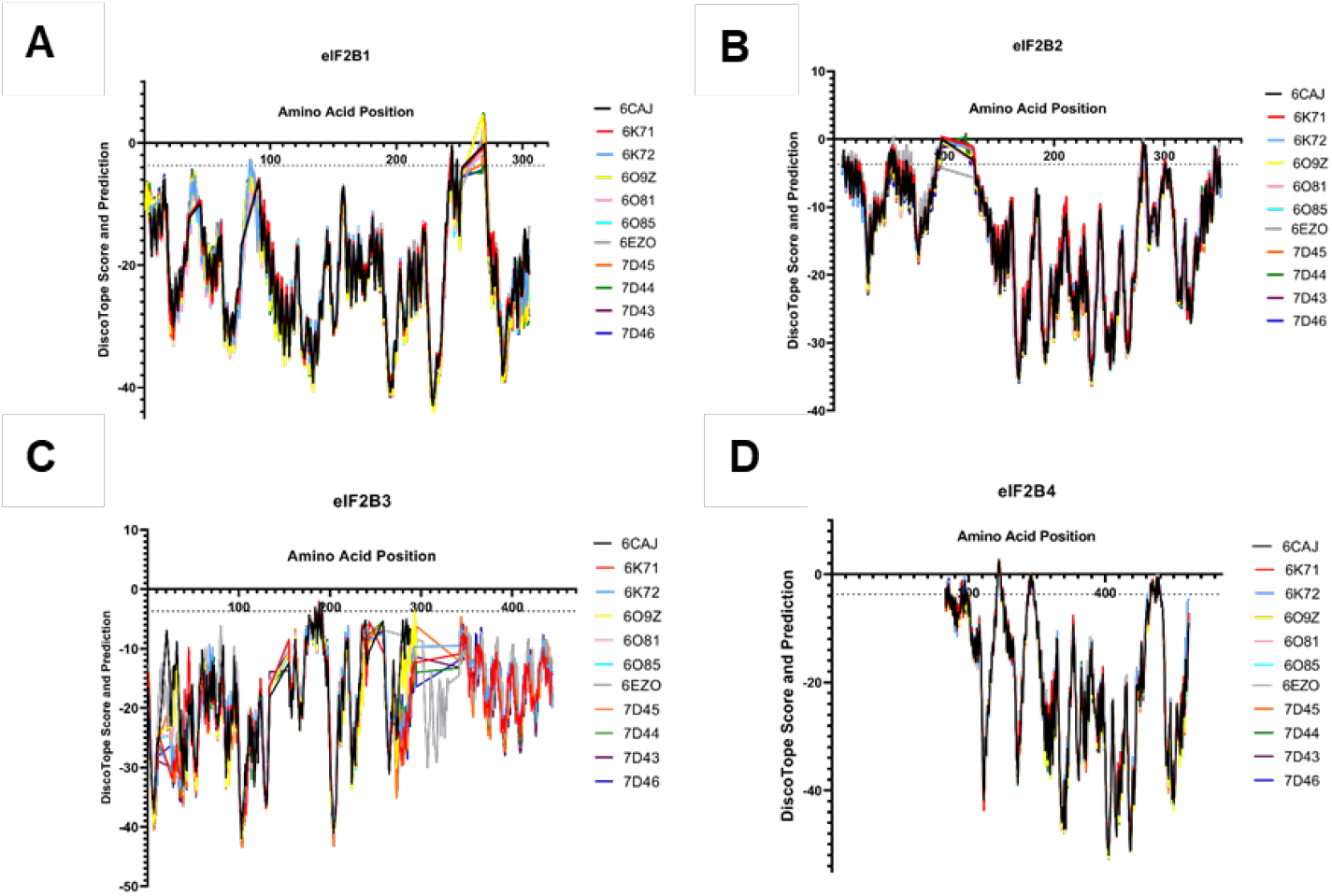
eIF2B epitope prediction using DiscoTope2.0. (A) DiscoTope2.0 Score with a ≥-3,7 threshold of eIF2Bα for each eIF2B PDB structure; (B) DiscoTope2.0 Score with a ≥-3,7 threshold of eIF2Bβ for each eIF2B PDB structure; (C) DiscoTope2.0 Score with a ≥-3,7 threshold of eIF2Bγ for each eIF2B PDB structure; (D) DiscoTope2.0 Score with a ≥-3,7 threshold of eIF2Bδ for each eIF2B PDB structure.

**Figure S4.**
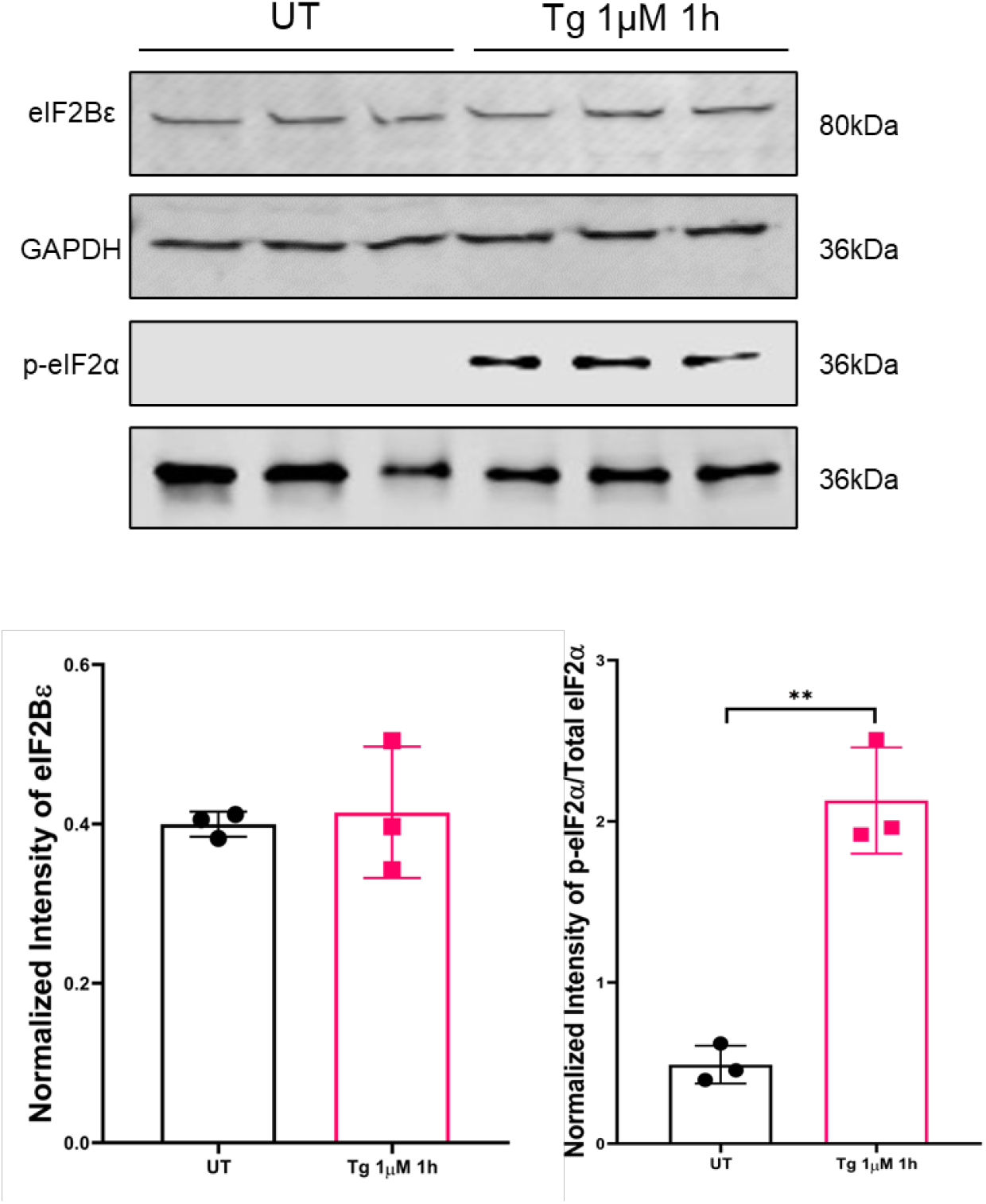
Expression levels of eIF2Bε following thapsigargin treatment. Western blot analysis of the level of eIF2Bε, p-eIF2α and total eIF2α expression in U373 cells either untreated or treated with 1μM Tg to induce cellular stress. Levels of eIF2Bε were normalized to levels of GAPDH (n=3). Levels of p-eIF2α were normalized to levels of total eIF2α (n=3). p values were derived from a Unpaired t-test, * *p ≤0.01

